# Phototaxis and the origin of visual eyes

**DOI:** 10.1101/027466

**Authors:** Nadine Randel, Gáspár Jékely

## Abstract

Vision allows animals to detect spatial differences in environmental light levels. High-resolution image-forming eyes evolved from low-resolution eyes via increases in photoreceptor cell number, improvements in optics and changes in the neural circuits that process spatially resolved photoreceptor input. However, the evolutionary origins of the first low-resolution visual systems have been unclear. We propose that the lowest-resolving (two-pixel) visual systems could initially have functioned in visual phototaxis. During visual phototaxis, such elementary visual systems compare light on either side of the body to regulate phototactic turns. Another, even simpler and non-visual strategy is characteristic of helical phototaxis, mediated by sensory-motor eyespots. The recent mapping of the complete neural circuitry (connectome) of an elementary visual system in the larva of the annelid *Platynereis dumerilii* sheds new light on the possible paths from non-visual to visual phototaxis and to image-forming vision. We outline an evolutionary scenario focusing on the neuronal circuitry to account for these transitions. We also present a comprehensive review of the structure of phototactic eyes in invertebrate larvae and assign them to the non-visual and visual categories. We propose that non-visual systems may have preceded visual phototactic systems in evolution that in turn may have repeatedly served as intermediates during the evolution of image-forming eyes.

## Introduction

Eyes are present in most animal phyla and show a great diversity of form and function (1,2). In the majority of phyla, eyes only mediate low-resolution vision or directional photoreception. Advanced high-resolution vision only evolved four times independently, in vertebrates, cephalopods, arthropods and alciopid polychaetes (3,4). This phyletic pattern suggests that the last common ancestor of bilaterians (referred to as “the urbilaterian”) did not possess complex visual eyes and may only have had simple eyes. The fossil record also suggests a humble beginning of eye evolution in the first bilaterians. For example, whereas many extant annelids have large visual eyes, the first annelids preserved as fossils from the lower and middle Cambrian lack evidence of eyes (5,6). Likewise, the earliest deuterostome and chordate fossils preserved animals with no morphologically distinguishable eyes, and only the first vertebrate fossils have eyes (7). Among the ecdysozoans, the lower Cambrian lobopodians (ancestral to euarthropods) only possessed simple bilateral eyespots (8). Unfortunately, it is not possible to infer from these adult fossils whether the larval stages possessed simple larval eyes. In contemporary animals, the larval stages can have simple eyes, even if the adults are eyeless, as for example in some annelids (9), brachiopods (10) or bryozoans (11).

Even if complex adult eyes were likely absent from early bilaterians, as suggested by the fossil record, the presence of photoreceptor cells and at least simple eyes is supported by the universal deployment of opsins as photopigments in eyes and the molecular similarities in eye development across bilaterians, as represented by well-studied model organisms. These similarities suggest that eyes as diverse as the vertebrate camera eye and the insect compound eye may have common evolutionary ancestry (12–15). This ancestral structure may have consisted of just a few photoreceptor cells and pigment cells, mediating directional photoreception or low-resolution image-forming vision. Eyes of such structural simplicity are widely distributed across metazoan phylogeny, and their detailed comparative study may hold the key to understanding the early steps of eye evolution.

One successful avenue for reconstructing the early steps of eye evolution has been the comparative study of photoreceptor cells (14). Based on morhological studies Eakin suggested that there are two major lines in the evolution of photoreceptors, one ciliary and the other rhabdomeric (16,17). The recognition that two major classes of opsins, the r-opsins and c-opsins, are associated with the rhabdomeric and ciliary photoreceptors, respectively, gave molecular support to Eakin’s theory (13). Comparative molecular studies focusing on the constituent cell types of eyes suggest that both rhabdomeric and ciliary photoreceptors (13), as well as pigment cells (18), may already have been present in the urbilaterian or even the stem eumetazoan (19). The deployment of similar transcriptional cascades across phyla may then reflect a conserved role in the specification of visual system primordia in the anterior nervous system and the differentiation of conserved cell types (20,21). These ancient cell types then further differentiated for the independent construction of complex eyes in different phyla. Significantly, most protostome eyes have rhabdomeric photoreceptors and vertebrate eyes have ciliary photoreceptors. However, derived rhabdomeric photoreceptors are likely still present in the vertebrate eye as melanopsin-containing retinal ganglion cells (20). Likewise, ciliary photoreceptors can be found in many protostomes, either as brain photoreceptors, or as parts of pigmented eyes (10,22,23) (Figure 1).

**Figure 1.**
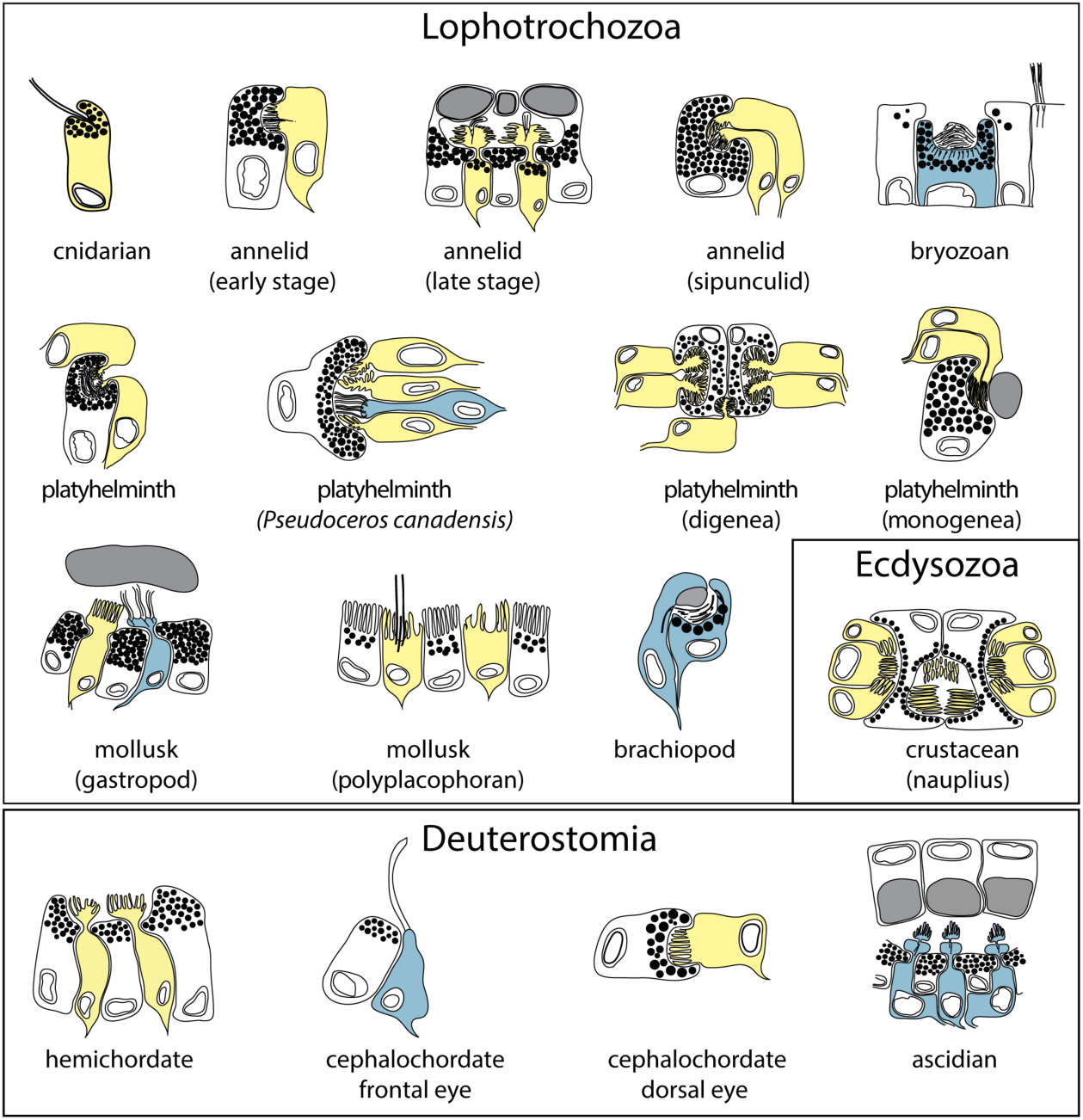
Diversity of simple eyes in planktonic larvae. Schematic drawings of simple eyes from marine invertebrate larvae. Rhabdomeric photoreceptors are shown in yellow, ciliary photoreceptors in blue, pigment cells in grey.

The distinction of rhabdomeric and ciliary photoreceptors and their corresponding opsins has been supported by many studies. It is to be noted, however, that opsins are also present in a large diversity of extraocular photoreceptors, cautioning that a one-opsin one-cell-type scenario may be too simplistic (24,25). The expression of opsins belonging to other opsin families, such as the Go-opsins and retinochromes in visual photoreceptors (26–28), further complicates the matter. In particular, a Go-opsin is expressed in ciliary photoreceptors in a mollusk, but its ortholog is expressed in rhabdomeric photoreceptors in an annelid (27,28). Another problem is that the molecular characteristics of photoreceptors in several animal groups have not yet been investigated, leaving open the question about the general validity of the rhabdomeric-ciliary distinction. However, if one focuses on photoreceptors with extended membrane surfaces found in pigmented eyes, the morphological and molecular distinction is well supported by the available data.

The history of other cell types of visual systems, including visual interneurons and motorneurons, is much less clear. One study suggested that visual interneurons are conserved across bilaterians, indicating that visual circuits may already have evolved along the bilaterian stem (29). However, despite recent progress (e.g. (30)), the development of visual interneurons is less well understood and detailed molecular studies have only been performed in a few species. One difficulty with the comparison of interneuron types across phyla, in contrast to photoreceptors, is the lack of easily identifiable conserved effector genes as cell-type markers. Nevertheless, further molecular and developmental comparisons may allow a fuller reconstruction of the cell-type diversity of eye circuits and other light-sensitive organ systems in the urbilaterian and the cnidarian-bilaterian common ancestor.

Here we propose a complementary approach, relying on the comparative study of the function and neural circuitry of simple larval eyes. We present a comprehensive classification of simple eyes of planktonic invertebrate larvae, based on morphological and functional criteria. We discuss the early evolution of vision from the functional and neural-circuit perspective and provide some guidelines for the experimental study of simple eyes.

## Simple eyes of planktonic larvae mediate phototaxis

Simple larval eyes (one to a few photoreceptor cells associated with shadowing pigment) are present in several species with planktonic dispersing larvae across multiple animal phyla (Figure 1 and Table 1). Despite their structural diversity, most eyes in marine invertebrate larvae likely mediate phototaxis. Phototaxis, defined as directional movement along a light vector towards (positive) or away (negative) from a light source, is widespread among marine larvae. Positive phototaxis is a common attribute of the early larval stages of animals with a pelagic-benthic life cycle. This behaviour contributes to upward migration in the water column and can facilitate larval dispersal. Older larval stages often become negatively phototactic and migrate towards the benthic zone shortly before larval settlement (31). In addition, many larvae have the capacity to switch between positive and negative phototaxis (mixed phototaxis). The ability to perform mixed phototaxis may help larvae to find a preferred water depth and stay in the water column for longer periods. Several environmental factors, including light intensity, UV-radiation, spectral distribution, temperature, salinity, chemicals, oxygen, food availability, and the presence of predators can switch the sign or modify the strength of phototaxis (32,33).

**Table 1.** Examples of the four major types of simple larval eyes we distinguished. The full list is in Supplementary Table 1. References: *Tripedalia cystopora* (76), *Autolytus prolifera* (50,77), *Capitella* sp. (78,79), *Harmothoe imbricata* (43,80), *Lanice conchilega* (50,81), *Neanthes succinea* (82), *Odontosyllis ctenostoma* (83), *Pectinaria koreni* (50), *Phyllodoce maculate* (84), *Phyllodoce mucosa* (85), *Platynereis dumerilii* (40,60,86), *Polygordius appendiculatus* (49,87), *Serpula vermicularis* (42,43), *Spirobranchus giganteus* (39,43,88,89), *Spirobranchus polycerus* (84,90,91), *Spirorbis spirorbis* (50), *Syllis amica* (83), *Golfingia misakiana* (92), Sipuncula larva (92), *Aporrhais pespelecani* (93), *Aporrhais sp.* (93), *Atlanta peroni* (94), *Bittium reticulatum* (55), *Carinaria lamarckii* (23), *Hermissenda crassicornis* (95), *Ilyanassa obsolete* (96,97), *Lacuna divaricata* (51), *Rostanga pulchra* (98), *Smaragdia sp*. (99), *Strombus sp.* (99), *Trinchesia aurantia* (100), *Katharina tunicata* (101), *Lepidochiton cinereus* (51), *Terebratalia transversa* (10), Loxosomatidae larva (56), *Notoplana alcinoi* (102), *Pseudoceros canadensis* (22), *Stylochus mediterraneus* (102,103), *Thysanozoon brocchii* (102), *Kronborgia isopodicola* (104), *Diplozoon paradoxum* (105), *Entobdells soleae* (106), *Euzetrema knoepffleri* (107), *Merizocotyle sp.* (108) *Neodiplorchis scaphipodis* (109), *Neoheterocotyle rhinobatidis* (110), *Polystoma integerrium* (111)*, Pseudodiplorchis americanus* (109), *Alloreadium lobatum* (112), *Fasciola hepatica* (112,113) *Heronimus chelidra* (112), *Heronimus mollis* (114), *Multicotyle purvisi* (115), *Philophtalmus megalurus* (116), *Spirorchis sp*. (112), *Bugula neritina* (57,117), *Bugula pacifica* (11), *Bugula simplex* (11), *Bugula stolonifera* (11,117,118), *Bugula turrita* (11), *Scrupocellaria bertholetti* (11,119), *Tricellaria occidentalis* (11,120), *Waterspora arcuata* (121), *Dactylopusia cf. tisboides* (52), *Balanus crenatus* (122), *Catalina sp*. (123), *Ptychodera flava* (123), *Glossobalanus marginatus* (124)*, Amaroucium constellatum* (125), *Ascidia nigra* (126), *Ciona intestinalis* (126–130), *Distaplia occidentalis* (128), *Phallusia mammillata* (126), *Branchiostoma floridae* (58,131,132).

| *Eye type* | Scanning photototactic eye | Visual phototactic eye |
| --- | --- | --- |
| *Photoreceptor type* |  |  |
| **Ciliary photoreceptor** | ascidian larval eyespot (*Ciona*); bryozoan larval eyespot | brachiopod larval cerebral eyes (*Terebratalia*); some mollusc veliger eyes |
| **Rhabdomeric photoreceptor** | box jellyfish larval eyespots; annelid trochophore eyespot (*Platynereis*); amphioxus early larval Hesse eyecup; nemertean larval eyes (?); polyplacophoran eyespot (?) | annelid nectochaete adult eye (*Platynereis*), some mollusc veliger eyes; crustacean nauplius larval eye; some platyhelminths; hemichordate late larva (?) |

Phototactic animals require a polarized body, at least one photoreceptor partially shaded by pigment, and a strategy to achieve spatial resolution (34). There are two fundamentally different mechanisms for this: scanning by helical rotation or head movement (non-visual scanning phototaxis) or spatial vision (visual phototaxis). These mechanisms are associated with two different types of eyes and neuronal systems; directional photoreceptors forming direct sensory-motor eyespots and more complex visual phototactic systems (Figure 2). We follow the definition of Nilsson (35) to distinguish between a directional photoreceptor that “relies on body movement to acquire information about the angular (spatial) distribution of light” and visual systems “with different photoreceptors pointing in different directions [to collect] spatial information… simultaneously without body movement.” In the following sections we discuss the function and anatomy of these two types of neural systems and how they mediate phototaxis.

**Figure 2.**
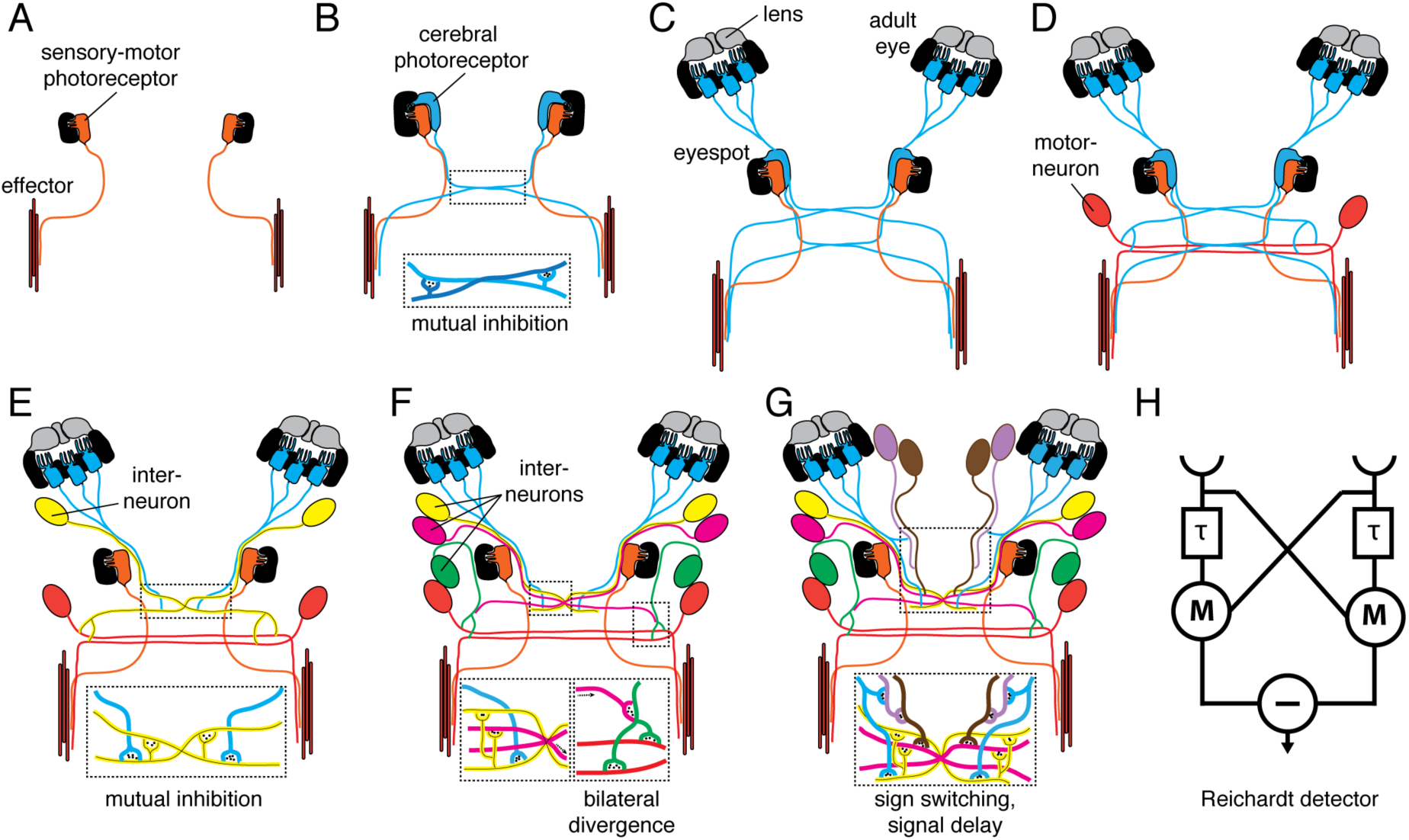
Transition scenario for the evolution of low-resolution visual eyes. Transition scenario for the evolution of low-resolution visual eyes from two-celled sensory-motor eyes. (A) Larval eyespot with a single photoreceptor cell (orange) mediates positive helical phototaxis by directly innervating an effector (muscle and ciliated cells are not distinguished for simplicity). (B) A duplication event leads to the development of a second photoreceptor cell (blue), which is able to mediate negative phototaxis. The development of mutual synaptic contacts between the contralateral photoreceptors allows contrast enhancement, representing the first step towards vision. (C) Relocation and duplication of the second photoreceptor cell (blue) results in the development of the cerebral eyes. The duplication of photoreceptors improves the signal to noise ratio, the development of a lens improves photon collection. (D) Integration of a motorneuron into the circuit and bilateral circuit divergence enables the signal to reach either the left or the right motor organ (effector). With this circuit, the animal is able to switch between positive and negative phototaxis, using the same eye. (E, F) Integration of one to several interneurons improves computational power and provides further possibilities for modulation (e.g. integration of other sensory inputs). (G) The development of a new type of primary interneuron (directly postsynaptic to the photoreceptor cells), which transmits the signal with a delay. This second primary interneuron might work together with the first primary interneuron (yellow) to form the first motion detector. (H) The Reichardt detector, a conceptual model for motion detection, adapted from (74). Red, sensory-motor photoreceptor of the eyespot and motorneurons; blue, adult eye photoreceptors; yellow, primary interneuron; pink, green, magenta, other interneurons.

## Non-visual scanning versus visual phototaxis

The simplest form of phototaxis is non-visual scanning phototaxis mediated by directional photoreceptors. It has two types, helical or conical scanning phototaxis, and horizontal scanning (yawing) phototaxis (36). Helical phototaxis is widespread among protists (not considered here, but see (34)) and metazoan larvae and relies on a direct sensory-motor coupling between a photoreceptor and an effector. Horizontal scanning phototaxis is found in some hydrozoan larvae (37,38) and relies on the left-right movement of the A-P body axis of the larva to scan the distribution of light. During non-visual phototaxis, the photoreceptor cell must be shaded from one side for directional sensing, and the pigment and the photoreceptor cell together form an eyespot or ocellus. Spatial information can only be obtained by rotating around the body axis during helical movement or by left-right bending of the A-P body axis. Such conical or horizontal scanning causes a temporal oscillation of the light stimulus. The magnitude of the oscillations depends on eyespot and body orientation and the spatial distribution of light (39). The eyespots are autonomous sensory-motor organs, and even if there are two or more of them, there is no informational exchange between them (40).

Helical phototaxis has been described in fine detail in the tadpole larva of the ascidian *Aplidium constellatum* (41) and the trochophore larva of the polychaete *Platynereis dumerilii* (40). In the ascidian tadpole, the eye signals to the muscle, and changes the direction of the helical swimming trajectory by tail bending (41). Although synaptic contacts have not yet been mapped, the navigation strategy ascidian larvae use is similar to that of the trochophore. The ascidian tadpole has only one eyespot, steering is thus only possible every 360 degrees of rotation. The *Platynereis* trochophore larva has two lateral eyespots, pointing in opposite directions due to the presence of shading pigment. The photoreceptor cell of each eyespot directly projects to and synapses on adjacent ciliated cells of the main locomotor ciliary band of the larva (prototroch). During rotational swimming along a helical axis, each eyespot receives an oscillating signal that repeats its peak value at every 360 degrees of rotation. An increase in light intensity activates the photoreceptor that signals to the ciliated cells, reducing ciliary beat frequency and changing the stroke pattern of cilia on the same body side. This leads to a small turn of the helical trajectory towards the light source. Since there are two eyes, steering episodes are repeated every 180 degrees of rotation until the larval trajectory is aligned with the light vector. The photoreceptor cells have to adapt quickly to be able to respond at every turn.

Even if there are two eyespots, as in many trochophores, the lack of a complex neural circuitry prevents the direct comparison of light inputs between the eyes on the two body sides. Rotational helical motion is therefore essential for phototaxis, as shown by computer simulations (40). The autonomy of the two eyespots can best be demonstrated by eye ablation experiments. In *Platynereis* trochophores, one eyespot is necessary and sufficient for phototaxis (40). Analogous to such one-eye-ablated larvae, some annelids in the early larval stage possess only a single eyespot and are phototactic (39,42,43). The early-stage cephalochordate larva also has a single eye, the Hesse eyecup (44), which may drive phototaxis. The young amphioxus larvae rotate around their body axis (45), suggesting that, at least in the early stages, phototaxis is of the helical non-visual type.

The neural mechanisms of non-visual horizontal scanning phototaxis in hydrozoan larvae are not known. Based on behavioural observations (38), it is likely that these larvae use directional photoreceptors and obtain spatial information by left-right bending along the A-P body axis. The photoreceptor cells may connect to myoepithelial cells in a simple sensory-motor circuit.

In contrast to non-visual phototaxis, visual phototaxis relies on the spatial comparison of light levels without body rotation or other scanning movements. Vision requires at least two photoreceptors pointing in different directions to detect differences in the spatial distribution of light (35). Visual systems also require a more complex underlying neural circuitry with the capacity for comparing light stimuli from the two or more photoreceptors.

The sensory-motor strategy and neuronal circuitry of visual phototaxis is best understood in the late stage (nectochaete) larvae of *Platynereis*. These larvae develop two pairs of eyes additional to the sensory-motor eyespots. These ‘adult eyes’ in the larva represent the developmental precursors to the adult’s visual eyes but with a larva-specific function. The four adult eyes and their neuronal circuitry (approximately 70 neurons) form a simple visual system that mediates late-larval visual phototaxis. During visual phototaxis, the larvae swim with cilia and steer with muscles (46). An imbalance in the light input to the two body sides triggers the contraction of the longitudinal muscles on or opposite to the side of illumination (depending on whether positive or negative phototaxis is taking place), leading to body bending during swimming. The coordinated action of at least one eye per body side is required for phototaxis, as demonstrated by eye ablations. If the adult eyes on one body side are ablated and the free-swimming larvae are exposed to non-directional light, they bend their body and swim in circles as long as they are illuminated. Contrary to sensory-motor eyespots,there is no adaptation to sustained illumination.

It is important to note that *Platynereis* nectochaete larvae do not obtain spatial information by left-right horizontal scanning, as the hydrozoan larvae, but by a visual system. The horizontal bending in *Platynereis* nectochaete larvae during visual phototaxis is a consequence of uneven illumination, and not a strategy to obtain information about uneven illumination. This visual strategy both works if the larva freely swims in water with helical turns or if it crawls on a surface on its ventral side. Nectochaete larvae can perform visual phototaxis in both helical and bilateral swimming mode.

The motor output of both non-visual and visual phototactic systems can be either locomotory cilia or muscles, or both. The motor organs used for propulsion versus steering may not necessarily be the same and in some animals the combination of the effectors used for phototaxis can change during larval development. For example, early stage *Platynereis* larvae swim and steer with cilia while late-stage larvae swim with cilia, but steer with muscles (40,46). Use of a combination of cilia- and muscle-based motor organs during swimming is reported in veliger larvae (mollusks) (47,48). In cephalochordates, muscles are used for fast response upon disturbance (45), and may play a role in turning during phototaxis in late-stage larvae.

## Classification of eye types in planktonic larvae

We performed a comprehensive survey of the literature and catalogued the available information (mostly ultrastructural) to build a framework for larval eye classification (Supplementary Table 1). Larval eyes are highly diverse in morphology, number and position. Based on the functional differences outlined above, we argue that the most important aspect of a classification should be whether the larval eyes and their neuronal circuitry constitute a visual system or not. We thus distinguish non-visual (helical or yawing scanning) phototacic eyes and visual phototactic eyes. The lack of functional studies precludes the assignment of some examples, but we can often distinguish the two major types based on the morphological data. Our classification also considers morphological criteria, such as the type of photoreceptor. Annelids (40,49,50), several mollusks (for example (51)), crustaceans (52), and some nemerteans (53) (although only juvenile and adult eyes have been described ultrastructurally (54)) have rhabdomeric eyespots. In contrast, brachiopods (10), some molluscs (55), entoprocts (56) and bryozoans (57) have ciliary eyespots, and can lack rhabdomeric photoreceptors. In some larvae both ciliary and rhabdomeric photoreceptors are present, and the two photoreceptor types may or may not be associated with pigment. In the annelids, rhabdomeric larval eyes are pigmented, but the brain ciliary photoreceptor cells are not associated with pigment cells (20). In the Müller’s larvae of marine flatworms the right eyespot contains rhabdomeric photoreceptors, while the left eyespot contains several rhabdomeric and one ciliary photoreceptor (22). The eye of some gastropod larvae undergoes ba transition from ciliary to rhabdomeric photoreceptor cells during development (23). Among the deuterostomes, both the ciliary and the rhabdomeric photoreceptors of cephalochordates are associated with melanin-containing pigment cells (58).

Another criterion for larval eye classification is the presence or absence of a lens. Some larval eyes have a lens that can form by the apical extension of the pigment cells, as in many annelids, by secretion by pigment cells and cornea cells or cornea cells alone, as in gastropods (51), or by lens cells, as in ascidians. The presence of a lens allows the larvae to collect more photons but is not correlated with the mode of phototaxis.

We also catalogued the number of cells in the eye and further morphological features, including the fine morphology of the sensory structures in the photoreceptor cells.

In order to classify eyes based on their neural circuitry, we used the innervation pattern of the photoreceptors and the type of effector (muscles or cilia), if known. Although the details of the innervation are only known in *Platynereis* and to some extent in the crustacean nauplius larva (52), for many taxa (annelids, mollusks, brachiopods, ectoprocts, hemichordates, bryozoa) the gross anatomy of photoreceptor cell projections is described (Supplementary Table 1). We also collected information about the habitat of the adult stages for most of the species.

The anatomy also allows inferences about function. Those eye photoreceptors that directly innervate the effectors, likely mediate helical phototaxis (e.g. bryozoan eyespots). Conversely, cerebral eyes that project to the brain neuropil or to the cerebral ganglion and thus connect to the effectors indirectly likely mediate visual responses (e.g. the eyes of veliger larvae). The known or inferred locomotor pattern (gliding on the surface with or without head movements versus helical or bilateral swimming) further helps to infer the type of phototaxis. We can use these criteria to assign several dozen examples across ten phyla to the categories of non-visual or visual phototactic eyes. We can extend this classification by photoreceptor type, to get a fourfold classification of simple larval eyes (Table 1). Further functional and neuronal connectome studies will be needed to confirm or modify our assignments.

In our framework, the most important functional distinction is between eyes that mediate non-visual phototaxis and those that mediate visual phototaxis. The most informative experiment for distinguishing a sensory-motor system from a visual system is the unilateral ablation of the eyes combined with phototaxis assays. If phototaxis still persists after the eye on one body side has been ablated, the larva must employ non-visual helical scanning phototaxis. An exception would be a vertebrate larva with high-resolution visual eyes (e.g. zebrafish), not considered here.

## A transition scenario for the origin of annelid visual eyes

Next, we would like to explore the possibility of an evolutionary transition from helical phototacic eyes to visual phototactic eyes. We use the example of annelid eyes, because this is the only case where the neuronal circuitry has been mapped for both eye types and where the behaviour has been described in detail (40,46). The connectome of the *Platynereis* larval visual system allows us to propose a detailed cell evolution scenario for the origin and evolution of annelid eyes. In this scenario, novel anatomies and functions arose from a common ancestor through duplication and divergence. Our model expands upon the ‘division of labour’ model for eye evolution (59), which did not provide an explanation for the origin of new functions. Many functions, such as visual contrast and bilateral signal-divergence, are absent from helical phototacic eyes and must have evolved *de novo*. Our scenario begins with an eyespot consisting of one pigment cell and one sensory-motor photoreceptor for mediating helical phototaxis. We posit that the first step in the evolution towards a visual system was the duplication of the photoreceptor cell to give rise to a new cell with a contralateral projection (Figure 2). The *Platynereis* trochophore larval eyespot represents a similar condition, containing one sensory-motor and one cerebral photoreceptor (60). The driving force behind this change may have been selection for the possibility of a developmental sign switch in phototaxis. The majority of marine invertebrate larvae are initially positively phototactic and develop negative phototaxis later in the life cycle. This functional switch is inherent to marine pelagic-benthic life cycles with a dispersing larva (31). One way to achieve this is to switch the neurotransmitter of the eye during development. There is pharmacological evidence suggesting that some bryozoan larvae use neurotransmitter change during development to attain sign switching (61). Our model in contrast proposes a rewiring step where a newly evolved photoreceptor innervated effector cilia on the contralateral side of the body. Such neuronal connections appearing at later developmental stages could have regulated negative phototaxis in late larvae. We posit that this newly evolved photoreceptor was the first ‘cerebral’ photoreceptor that had to cross the neuronal midline (not necessarily the first neuron). The crossing of the axons of the first cerebral photoreceptors from the two body sides opened up the possibility for bilateral communication. One solution for this would be for the two cells to evolve mutual inhibitory connections, representing a simple motif for contrast enhancement. Such mutual connections may have been the first steps towards the evolution of visual phototaxis. Cerebral photoreceptors crossing the midline and communicating with each other could have provided the ability to compare light inputs between the two sides of the body. The functional elegance of such a system lies in its property to detect contrast, independent of total light intensity. This is because the level of activation of an eye is a function of the incident light minus the signal from the other eye. This represents an evolutionary stage where the sensory-motor eyes already form a visual circuit.

Such a system would also have required the tuning of the adaptation kinetics of the photoreceptors. Visual comparisons during phototaxis cannot work if the two eyes constantly and independently dark-adapt and light-adapt. This would make any comparison difficult. The adult eyes that form a visual phototactic system in the *Platynereis* larva, can maintain their activation for tens of seconds when illuminated.

A possible further step could have been the duplication and dorsal migration of the cerebral photoreceptors to give rise to the annelid adult eyes. These eyes may have initially connected to the contralateral muscles and/or ciliary bands. In our scenario the mutual inhibitory connections are maintained between the adult eye photoreceptors to enhance visual contrast.

The next step in our model for the evolution of a visual system is the evolution of motorneurons.

Motorneurons evolved either via the duplication of the photoreceptors or were recruited to the eye circuit from pre-existing motorneurons that already regulated turning behavior or ciliary activity for example in a haptic circuit. At this stage, the photoreceptors could have lost direct contact to the effector cells, to contact them indirectly via the motorneurons. The introduction of motorneurons to the circuit decoupled sensory from motor functions.

Further duplications of the photoreceptors may have given rise to the interneurons of the primary optic neuropil of the annelid visual system. One subset of these, the primary interneurons, retained the mutual inhibitory connections and received direct input from the photoreceptors. The connection of the interneurons to both contralateral and ipsilateral motorneurons may have allowed the bilateral divergence of the light signal, allowing more complex modulation of turning decision and the ability to rapidly switch between positive and negative phototaxis (Figure 2). The number of photoreceptors also increased, increasing the signal-to-noise ratio by averaging signals from several photoreceptors onto a single interneuron. This situation can be seen in *Platynereis*, where several photoreceptors synapse on the same primary interneuron, a circuit motif enabling signal averaging. The evolution of a lens in the adult eye increased the number of photons captured, increasing sensitivity.

Any such scenario is conjectural and contentious. However, we think that it is important to propose specific theories to stimulate further thinking about circuit evolution. There are some objections that one could raise. For example, contralateral projections may have already been present before the evolution of eyes, for example in haptic or chemosensory circuits. Different sensory systems in animals may have a common origin (62), suggesting that the first eye circuits may have evolved on a pre-existing scaffold of haptic or other systems. However, these different sensory modalities may already have diversified in radial ancestors, where contralateral projections did not exist.

## Were visual phototactic eyes present in the urbilaterian?

The basic logic of the annelid scenario may be applied to the evolution of eyes in general. If simple pigmented eyes trace back to the stem bilaterian, as is likely (13), the succession of events we inferred from the state of extant annelid eyes may have happened before the annelid stem, possibly in the lophotrochozoan stem (sensu (63)) or even deeper. Currently there is insufficient comparative evidence to decide. Among the lophotrochozoans, the eyespots of annelid, mollusk, nemertean and polyclad flatworm larvae may be homologous, since they commonly derive from the first-quartet micromere derivatives of the A and C quadrants (1a, 1c) during spiral cleavage (64). An exception is the larval eyespot of chitons (65). The comparison of visual eyes is less straightforward, and some investigators for example consider the adult eyes of annelids and mollusks non-homologous (66). Alternatively, annelid, mollusc and nemertean pigment-cup eyes may be homologous (13).

Instead of tracing homologies, here we argue based on strong functional constraints that simple visual eyes may have already appeared in stem bilaterians. We argue that the advent of a bilateral body plan and bilateral locomotion could have profoundly affected the evolution of eyes. Bilateral symmetry and locomotion evolved along the bilaterian stem lineage from presumably radial ancestors (67). However, many bilaterians display helical locomotion as larvae, with fundamental importance for phototactic sensory-motor processing. Stem bilaterians may have had a dispersing larval stage (68–70), propagating via helical ciliary swimming, and an adult stage that used bilateral locomotion during crawling on surfaces.

We propose that the radial-to-bilateral transition in body plan and helical-to-bilateral transition in locomotor pattern may have driven the evolution of visual phototaxis from helical phototaxis. Helical phototaxis simply cannot work if the locomotion pattern is bilateral (but visual phototaxis can work during helical motion). Annelid early-stage and late-stage larvae illustrate this nicely. The trochophore larvae use helical swimming and sensory-motor eyes. However, nectochaete larvae can both rotate around their A-P axis when swimming or crawl on a surface without axial rotation and perform visual phototaxis using the adult eyes under both modes of locomotion. The implication of these functional constraints is that if the urbilaterian had eyes and bilateral locomotion in post-larval stages, these eyes very likely constituted a visual system. Again, as outlined above in detail for the annelids, the most likely origin of such a visual system is the sensory-motor eyes mediating phototaxis in the helically swimming larval stage. The sensory-motor eyes in bilaterian larvae could have originated from similar larval eyes in radial ancestors. One extant example is represented by the larval phototactic eyespots of the box jellyfish *Tripedalia cystopora* (76). As long as phototaxis and larval locomotion are helical, there is no major theoretical difference between a radial and a bilateral larva regarding the strategy of phototaxis and its neural requirements.

## Evolution of image-forming eyes from phototactic visual eyes

We suggest that image-forming eyes in Bilateria may have evolved multiple times (vertebrates, annelids, mollusks) from visual phototactic eyes. How could a visual phototactic system evolve into a low-resolution image-forming eye? Image-forming eyes have several properties not found in phototactic eyes, including motion detection, optokinetic response (e.g. to detect a drift in body position in a current), approach-detection (predator detection) or prey following. These behaviours all require several photoreceptors and complicated neuronal circuitry (71). However, phototactic visual eyes have several features that represent intermediates for more complicated circuits. For example, image-forming eyes have spatial resolution within the same eye. Eyes mediating visual phototaxis, such as the adult eyes in the *Platynereis* larva, have no resolving power and spatial resolution derives from the use of multiple eyes for a visual task (46). However, these eyes, constituting elementary visual systems, can contain multiple photoreceptors that may have initially evolved for signal averaging, but could have served as a substrate for structural diversification. A lens is also present in many phototactic visual eyes. Any lens is better than no lens, since lenses allow the eye to collect more photons. This could already have improved the sensitivity of phototactic visual eyes, and paved the way for using the lens for a more advanced focusing function.

Another key property of visual eyes is motion detection. Motion detection is a prerequisite for approach selectivity, optokinetic response, or prey following. Relatively simple circuits can have the ability to detect motion, including the Barlow-Levick detector (72) and the Reichardt detector (73,74). Both of these motion detection circuits have some shared basic properties. They require that the photoreceptors have directional sensitivity, a property already enabled by the pigment cup and lens of the phototactic visual eye. Signals from the photoreceptors then converge on the same interneuron, but signals from one photoreceptor are delayed (75). The interneuron is only active if signals from the two photoreceptors coincide. Such a circuit can in principle evolve by intercalating one interneuron between the integrator interneuron and the photoreceptors. In the *Platynereis* visual system, several interneuron types with an unknown function were described (46).

These may develop into the more complicated image-forming circuitry of the adult worm.

Discovering the details of neural circuit evolution for such visual tasks will require the mapping and analysis of complete circuits for many phototactic visual systems and image-forming eyes. The expectation is that through the study of different developmental stages in the same animal, as well as different species displaying varying levels of visual acuity, fine gradations will be discovered. Eventually, comparative connectomics may allow the reconstruction of evolutionary paths linking the simplest eyespots to more complex high-resolution visual eyes. We anticipate that the gradations will be fine enough that even Darwin would “conquer the cold shudder” he felt when thinking about the evolution of the eye.

## Conclusions

We proposed a functional classification of simple eyes in planktonic larvae that could serve as a guiding principle for future functional and anatomical studies. Further studies of simple eyes across a broader sampling of metazoans should lead to a more reliable reconstruction of ancestral states at distinct deep nodes of the metazoan phylogeny. Clearly, an integrative approach, combining the molecular, functional and neural network perspectives, will be needed in order to obtain increasingly sharper pictures of early eye evolution.

## Additional Information

## Acknowledgments

We thank Elizabeth Williams for comments on the manuscript.

## Authors’ Contributions

NR compiled Table 1 based on the literature. NR and GJ wrote the paper.

---

Supplementary Table 1. Morphological characteristics of simple larval eyes.

The table is based on an extensive survey of the literature on larval eye ultrastructure and function. The data on the adult’s habitat and location are based on the Materials and Methods sections of the cited papers describing the collection of the studied specimen. Note that the larval stages are always planktonic. Abbreviations: ae, adult eye; ci, ciliary photoreceptor; i/e, inverse or everse type of eye; le, larval eye; rh, rhabdomeric photoreceptor, PRC, photoreceptor cell; pt, sign of phototaxis.

